# Septin function tunes lipid kinase activity and phosphatidylinositol 4,5 bisphosphate turnover during G-protein coupled PLC signaling *in vivo*

**DOI:** 10.1101/2021.05.14.444211

**Authors:** Aastha Kumari, Avishek Ghosh, Sourav Kolay, Padinjat Raghu

## Abstract

The hydrolysis of phosphatidylinositol 4,5− bisphosphate [PI(4,5)P_2_] at the plasma membrane by receptor activated phospholipase C (PLC) activity is a conserved mechanism of signal transduction. Given the low abundance of PI(4,5)P_2_ at the plasma membrane, its hydrolysis needs to be coupled to lipid resynthesis to ensure continued PLC activity during receptor activation. However, the mechanism by which PI(4,5)P_2_ depletion during signalling is coupled to its resynthesis remains unknown. PI(4,5)P_2_ synthesis is catalyzed by lipid kinase activity and the phosphorylation of phosphatidylinositol 4 phosphate (PI4P) by phosphatidylinositol 4 phosphate 5 kinase (PIP5K) is the final step in this process. In *Drosophila* photoreceptors, sensory transduction of photon absorption is transduced into PLC activity leading to an electrical response to light. During this process, PI(4,5)P_2_ is resynthesized by a PIP5K activity but the mechanism by which the activity of this enzyme is coupled to PLC signalling is not known. In this study, we identify a unique protein isoform of dPIP5K, dPIP5K^L^ that is both necessary and sufficient to mediate PI(4,5)P_2_ synthesis during phototransduction. The activity of dPIP5K^L^ *in vitro* is enhanced by depletion of PNUT, a non-redundant subunit of the septin family of GTP binding proteins and *in vivo*, depletion of *pnut* rescues the effect of dPIP5K^L^ depletion on the light response and PI(4,5)P_2_ resynthesis during PLC signalling. Lastly we find that depletion of Septin Interacting Protein 1 (Sip1), previously shown to bind PNUT, phenocopies the effect of *dPIP5K^L^* depletion *in vivo*. Thus, our work defines a septin 7 and Sip1 mediated mechanism through which PIP5K activity is coupled to ongoing PLC mediated PI(4,5)P_2_ depletion.

## Introduction

In many eukaryotic cells the binding of extracellular ligands to plasma membrane receptors is transduced by the activation of phospholipase C (PLC). PLC hydrolyzes the membrane lipid phosphatidylinositol 4, 5− bisphosphate [PI(4,5)P_2_] on the inner leaflet of the plasma membrane to generate inositol 1,4, 5 trisphosphate (IP_3_) and diacylglycerol (DAG), both of which encode information leading to cellular responses (Berridge, 2009). PI(4,5)P_2_ is present in low amounts at the plasma membrane and during the high rates of PLC activation seen during cell surface receptor activation, PI(4,5)P_2_ at the plasma membrane would be rapidly depleted leading to loss of signalling sensitivity. However, PI(4,5)P_2_ levels at the plasma membrane are relatively stable and the depletion of this lipid during cell PLC activation is typically only transient. Thus, the utilization of PI(4,5)P_2_ by PLC is coupled to the synthesis of this lipid; in eukaryotes, the principal route of PI(4,5)P_2_ synthesis is through the sequential phosphorylation of phosphatidylinositol (PI) at positions 4 and 5 by phosphatidylinositol 4 kinase (Balakrishnan et al., 2018; Nakatsu et al., 2012) and phosphatidylinositol 4 phosphate 5-kinase (Chakrabarti et al., 2015; Stephens et al., 1991)[reviewed in (Kolay et al., 2016)]. However, in order for these enzymes to resynthesize PI(4,5)P_2_ in response to its utilization by PLC activity, it is necessary that plasma membrane PI(4,5)P_2_ levels are sensed and any drop in the levels of this lipid be communicated to the enzymatic machinery that regulates its synthesis.

*Drosophila* photoreceptors are an influential model for the analysis of G-protein coupled PLC signalling (Raghu et al., 2012). In these cells, the absorption of light by rhodopsin in photoreceptors is transduced into ion channel activity through G-protein coupled PLCβ activation leading to the hydrolysis of PI(4,5)P_2_. As part of their visual ecology, *Drosophila* photoreceptors experience bright light illumination leading to high rates of PLC activation (Fain et al., 2010); despite this, fly photoreceptors do not undergo rapid inactivation and must therefore have mechanisms to sustain PI(4,5)P_2_ levels during high rates of PLC β activity experienced in bright light illumination. Previous studies have shown that in photoreceptors, during illumination, PI(4,5)P_2_ resynthesis is regulated by a number of molecules including the phosphatidylinositol transfer protein RDGB (Yadav et al., 2015), PI4KIIIα that generates PI4P (Balakrishnan et al., 2018) and a PIP5K (dPIP5K) that phosphorylates PI4P to generate a pool of PI(4,5)P_2_ that is specifically required for normal phototransduction (Chakrabarti et al., 2015). However, the mechanism by which dPIP5K activity is regulated during light induced PLC activity in photoreceptors remains unknown.

Septins are a group of evolutionarily conserved, GTP binding proteins first discovered in yeast but now known to be widely distributed in the animal kingdom (Cao et al., 2009). Septins are part of a multiprotein complex composed of four classes of subunits; the complex is assembled at the inner leaflet of the plasma membrane using monomers and each class of monomer is composed of multiple members (Fung et al., 2014). They have been implicated in many processes including cell polarity, mitotic spindle positioning and cytokinesis. At the sub-cellular level, septins have been implicated in cytoskeletal organization (Mostowy and Cossart, 2012) through the formation of polymeric, filamentous structures. They have also been studied in the context of plasma membrane compartmentalization through a barrier function and a role for septins in regulating store operated calcium influx at the plasma membrane has also been reported (Sharma et al., 2013). In animals, the septin gene family is expanded to encode upto thirteen members (Nishihama et al., 2011) divided into four classes and septin oligomers usually include members from each class. In addition to protein interactions between the monomers, septins have also been shown to interact with many cellular proteins and through this mechanism regulate sub-cellular processes (Neubauer and Zieger, 2017). In organisms such as *Drosophila*, the septin gene family is less diverse; for example the Sept7 subgroup is encoded by a single gene, *pnut* (Neufeld and Rubin, 1994). Null mutations in *pnut* are homozygous lethal during development; however clonal analysis as well as studies of *pnut/+* in *Drosophila* have implicated *pnut* function in cytoskeletal regulation of cell shape and development in many tissues (Adam et al., 2000; Menon and Gaestel, 2015) and also in the regulation of store operated calcium influx (Deb et al., 2016). Importantly, septin monomers have been shown to bind PI(4,5)P_2_ (Zhang et al., 1999) through an N-terminal phosphoinositide binding domain. This PI(4,5)P_2_ binding has been shown to modulate the activity of this class of proteins (Bertin et al., 2010; Tanaka-Takiguchi et al., 2009). Thus, septins are a class of proteins localized to the inner leaflet of the plasma membrane and bind PI(4,5)P_2_. Given that PLC activity also hydrolyzes PI(4,5)P_2_ at the inner leaflet of the plasma membrane, septins are suitable candidates for proteins sensors that can detect changes in PI(4,5)P_2_ levels. However, to date there is no report of the regulation of PI(4,5)P_2_ levels at the plasma membrane by septins.

In this study we identify an isoform of dPIP5K-dPIP5K^L^, that is enriched in the eye which is both necessary and sufficient to regulate PI(4,5)P_2_ synthesis during PLC signalling to generate a normal response to light. *pnut* depletion enhances the activity of dPIP5K^L^ *in vitro* and also enhances dPIP5K^L^ mediated synthesis of PI(4,5)P_2_ during PLC signalling in photoreceptors. Further, we find that ***<u>S</u>***eptin ***<u>i</u>***nteracting ***<u>p</u>***rotein 1 (Sip1) is required for dPIP5K^L^ mediated PI(4,5)P_2_ synthesis in photoreceptors. Together our findings define PNUT/Sip1 as a regulator of PI(4,5)P_2_ synthesis during G-protein coupled PLC signalling *in vivo*.

## Results

### Identification of dPIP5K^L^, a novel eye enriched isoform of dPIP5K

dPIP5K is an eye enriched PIP5K which produces PI(4,5)P_2_ required for normal phototransduction (Chakrabarti et al., 2015). In a loss of function mutant of *dPIP5K* (*dPIP5K^18^*) the electrical response to light is severely reduced. We observed that the reduced light response could be rescued by reconstituting all the isoform of dPIP5K (*dPIP5K^BAC^)*, however reconstitution of a single isoform (*dPIP5K^S^*) was unable to restore the light response (Fig 1A). Based on the differential rescue and given that the *dPIP5K* gene is alternatively spliced, it seemed possible that a splice variant other than *dPIP5K^S^* must be required support PIP_2_ resynthesis during phototransduction. To identify such an isoform, we reverse transcribed total RNA extracted from 1-day old adult fly retinae using gene specific primers, designed based on the genomic sequence of the *dPIP5K* locus. Sequence analysis of the amplified cDNA product revealed a novel isoform (Full sequence in Fig S1), distinct from the already known isoform *dPIP5K^S^*. This transcript encoded a unique C-terminus of 273 amino acids previously not reported for existing isoforms. Therefore, we named this longer isoform as *dPIP5K^L^* and the known shorter isoform as *dPIP5K^S^* (Fig 1B,Fig S1). This new *dPIP5K^L^* splice variant encoded a protein isoform with a unique, extended C terminal domain (Fig 1B). Through a comparison of RNA extracted from wild type and *so^D^* mutant (without eyes) heads, we also found that the *dPIP5K^L^* transcript is enriched in the eye (Fig 1D). Protein blots from *Drosophila* S2R+ cells transfected with *dPIP5K^L^* shows the expression of a protein with the predicted molecular mass of 100kDa (Fig 1E). Previously we have reported that dPIP5K is localized to the microvillar plasma membrane, using an anti-dPIP5K antibody **[3]**. That antibody was raised against a common region of both dPIP5K^L^ and dPIP5K^S^ and hence could not be used to detect specific isoforms by immunofluorescence. To detect the localization of each isoform, we generated epitope-tagged constructs for each of them (dPIP5K^S^∷HA and dPIP5K^L^∷GFP). When reconstituted individually into *dPIP5K^18^* (a protein null allele) photoreceptors, we found that dPIP5K^L^ localizes to the microvillar plasma membrane (Fig 1F), whereas dPIP5K^S^ was distributed throughout the cell body of photoreceptors (Fig 1G). Thus, overall dPIP5K^L^ is an eye enriched isoform of dPIP5K, localized to the microvillar membrane. These observations prompted us to study if dPIP5K^L^, with its unique C terminal domain could be relevant to phototransduction.

**Figure 1:**
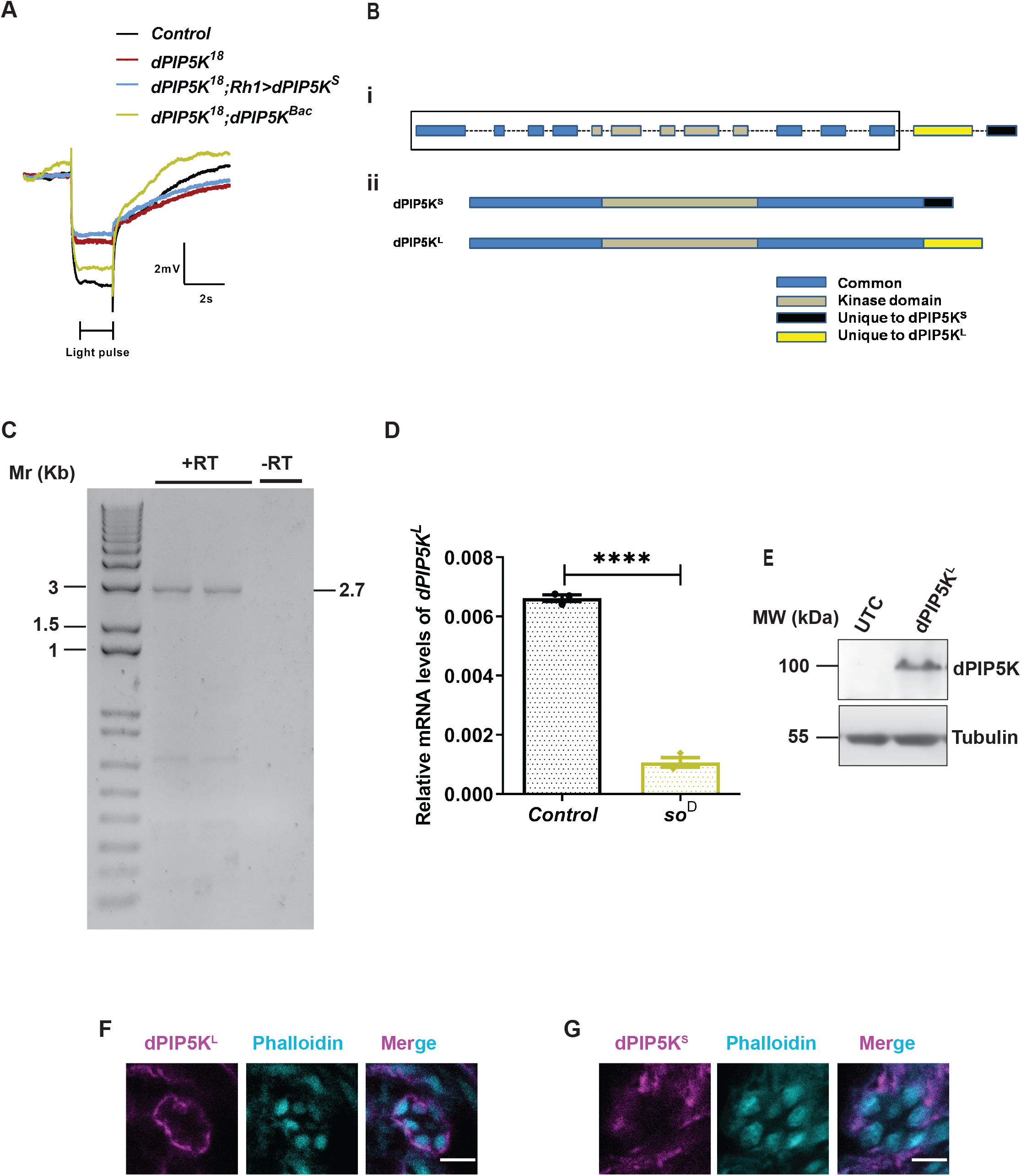
dPIP5K^L^ is an eye enriched isoform of *dPIP5K*. **(A)** Representative ERG trace from 1-day old flies of the indicated genotypes. The duration of the stimulus-a flash of green light is shown. Y axis shows response amplitude (mV), X axis indicates duration of the recording(s). **(B)** (i) Schematic illustrating the genomic structure of the dPIP5K gene. Exons are represented as rectangles and dashed lines mark the introns. The region common to both isoforms is marked out by the black rectangular box (ii) The protein structure of the two isoforms: dPIP5K^S^ and dPIP5K^L^ is shown. Domains that are identical between both isoforms are marked in blue, grey marks the common kinase domain, the C terminal region of dPIP5K^S^ is marked in black (132 amino acids) and the C terminal of dPIP5K^L^ is marked in yellow (273 amino acids) **(C)** Agarose gel image showing the amplification of cDNA for *dPIP5K^L^* at 2.7 Kb. +RT and –RT represent presence or absence of the reverse transcriptase enzyme in the reaction mix respectively. Amplification of the cDNA is seen only in the +RT reactions (loaded in duplicates). **(D)** Quantitative reverse transcription-PCR from RNA extracted from the heads of 1− day old flies showing enrichment of *dPIP5K^L^* transcript in wild type heads over *so^D^*. Y axis represents relative transcript level for *dPIP5K^L^*, normalized to internal control (RP49: ribosomal protein 49). X axis represents the genotypes (n=3 biological replicates). Error bars represent S.E.M. **(E)** Western blot from S2R+ cell extract prepared after transfecting cells with *dPIP5K^L^*, probed with anti-dPIP5K antibody(Chakrabarti et al., 2015). A 100 kDa band corresponding to dPIP5K^L^ is observed in transfected cells. Tubulin at 55kDa is used as loading control. (F,G) Confocal images of retinae from 1- day old transgenic flies over expressing dPIP5K^L^∷GFP **(F)**and dPIP5K^S^ ∷HA **(G)**in *dPIP5K^18^* background. Phalloidin (cyan) marks the location of the rhabdomere in each photoreceptor. dPIP5K^L^ is detected using anti GFP antibody (magenta) and dPIP5K^S^ is detected using anti HA antibody (magenta). Scale bar = 5μm. Data information: XY plots and scatter dot plots with mean±S.E.M are shown. Statistical tests: (D) two tailed unpaired t test with Welch correction. ****p<0.0001.

### *dPIP5K^L^* encodes a polypeptide with PIP5K activity *in vitro*

To understand the functions of dPIP5K^L^and dPIP5K^S^, we measured their enzymatic activities. PIP5K enzymes use PI4P as a substrate and generate PI(4,5)P_2_. *Drosophila* S2R+ cells do not express dPIP5K. For the enzymatic assay, we used the cell lysate from *Drosophila* S2R+ cells transiently expressing either dPIP5K^L^ or dPIP5K^S^ (Fig 2A). Using these lysates, we set up an *in vitro* kinase assay to quantify the extent of PIP_2_ produced by the conversion of the exogenously supplied substrate 37:4 PI4P to PI(4,5)P_2_. The synthetic substrate and product, of unique molecular mass was detected and quantified by LC-MS/MS. The extent of enzymatic conversion was measured by the response ratio which is derived by dividing the area under the curve (AUC) of synthetic PI(4,5)P_2_ signal by the AUC of synthetic PI4P signal. Under these *in vitro* assay conditions, we observed that the response ratio of *dPIP5K^L^* transfected lysates were significantly higher than that from untransfected cell lysates, signifying a high PIP5K activity. (Fig 2B). Also, dPIP5K^S^ was catalytically inactive *in vitro;* the response ratio was not different between *dPIP5K^S^* transfected cell lysates and those from untransfected controls (Fig 2B). The baseline PIP5K activity seen in untransfected cell lysates most likely arises from the endogenous PIP5K expressed in these cells encoded by *sktl* (Hassan et al., 1998), the only other gene in the *Drosophila* genome that encodes a PIP5K enzyme. We also confirmed that the PIP5K activity of dPIP5K^L^ is due to its kinase domain as a catalytically dead version of dPIP5K^L^ (dPIP5K^L-K189A^) did not elicit a significantly different response ratio compared to untransfected controls (Fig 2C, D). Thus, we conclude that dPIP5K^L^ has a high PIP5K activity *in vitro* whereas dPIP5K^S^ by itself does not exhibit PIP5K activity.

**Figure 2:**
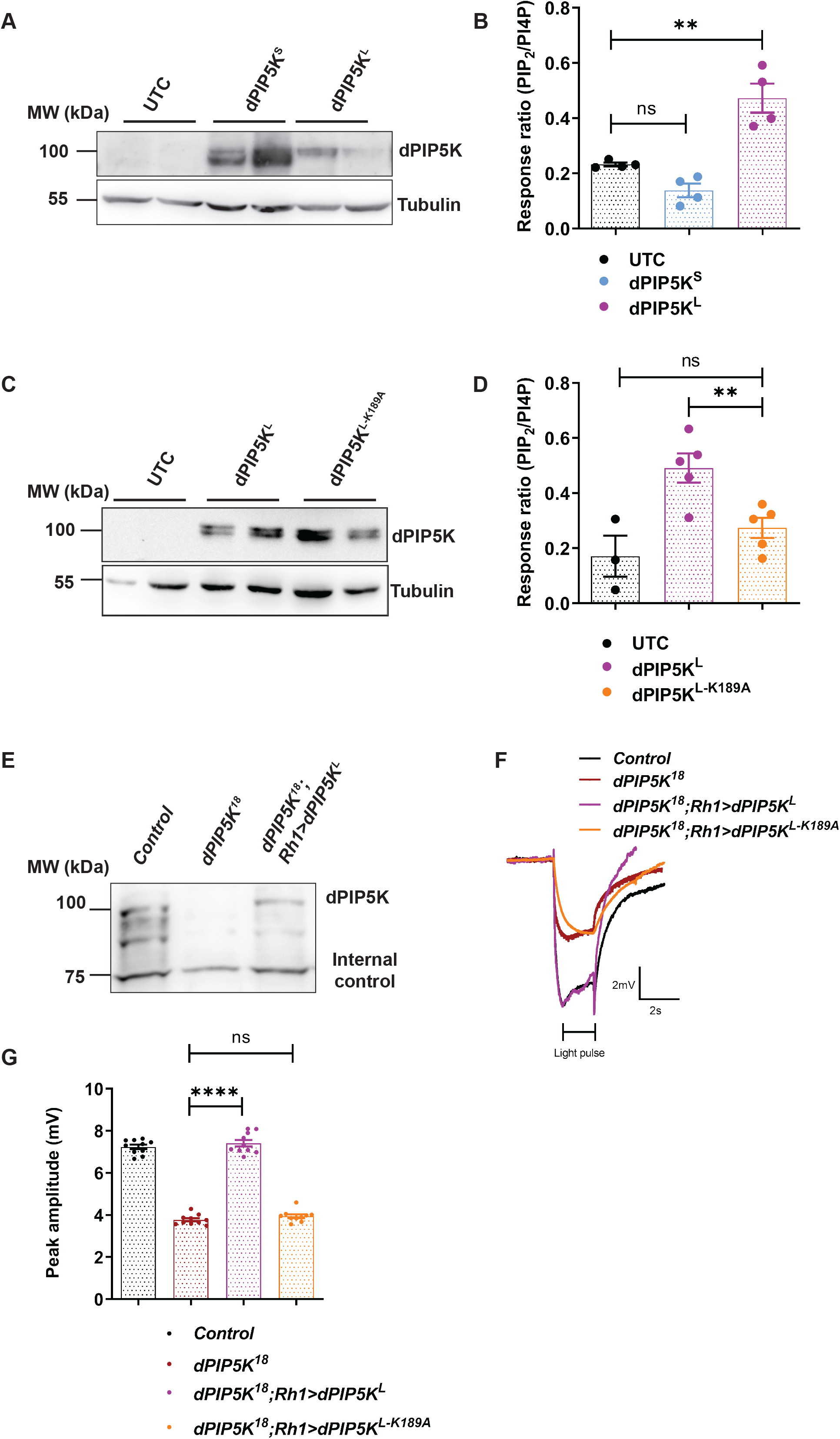
dPIP5K^L^shows PIP5K activity *in vitro* and is sufficient to sustain phototransduction *in vivo*. **(A)** Representative western blot showing expression of dPIP5K^L^ and dPIP5K^S^ from S2R+ cells extract transfected with the respective constructs. Samples are loaded in duplicates and probed with dPIP5K antibody. Tubulin at 55kDa is used as loading control. **(B)** Response ratio of product:substrate [37:4-PIP_2_:37:4-PI4P] in an *in vitro* kinase assay with cell free lysate prepared from S2R+ cells transfected with *dPIP5K^L^* or *dPIP5K^S^* (n=4 biological replicates). This graph is plotted by pooling data from 2 experiments. Error bars represent S.E.M. **(C)** Representative western blot showing expression of dPIP5K^L^ and dPIP5K^L-K189A^ from S2R+ cells extract transfected with the respective constructs. Samples are loaded in duplicates and probed with dPIP5K antibody. Tubulin at 55kDa is used as loading control. **(D)** Response ratio of product:substrate [37:4-PIP_2_:37:4-PI4P] in an *in vitro* kinase assay with cell free lysate prepared from S2R+ cells transfected with *dPIP5K^L^* or *dPIP5K^L-K189A^* (n=3 biological replicates for UTC and n=5 for dPIP5K^L^ and dPIP5K^LK189A^). Error bars represent S.E.M. **(E)** Western blot from 1-day old fly retinae showing expression of dPIP5K^L^ in *dPIP5K^18^* background (100kDa band). Internal control is shown at 75kDa. **(F)** Representative ERG trace from 1- day old flies of the indicated genotypes. The duration of the stimulus-a 2s flash of green light is shown. Y axis shows ERG amplitude (mV), X axis shows duration of the recording (s). **(G)** Quantification of the peak amplitude for ERG response in each of the mentioned genotypes. Y axis represents peak amplitude and X axis shows the genotypes. Each data point represents an individual fly tested (n=10 flies). Error bars represent S.E.M. Data information: XY plots and scatter dot plots with mean±S.E.M are shown. Statistical tests: (B,D,G) One way Anova with post hoc Tukey’s multiple pairwise comparison.ns-Not significant; *—p<0.01; ****p<0.0001

### dPIP5K^L^ is sufficient to sustain phototransduction

In order to study the functional relevance of dPIP5K^L^,we tested if it could rescue the ERG defects seen in *dPIP5K^18^*. We expressed dPIP5K^L^ in *dPIP5K^18^*. We observed that most of the flies eclosed with eyes having morphological defects, hence could not be used for ERG recordings; this phenotype is likely due to altered developmental events and was dependent on the kinase activity of dPIP5K^L^ as it was not seen on expression of dPIP5K^L-K189A^ (Fig S2A). However, on rearing the cross at 18°C during development, we could obtain some escapers with normal eye morphology. Western blot analysis of lysates from the eyes of such animals showed the selective re-expression of the dPIP5K^L^ isoform (Fig 2E). ERG recordings from these flies showed that expressing dPIP5K^L^ alone in *dPIP5K^18^* flies completely rescues both the amplitude as well as transients in the ERG (Fig 2F). We also observed that when expressed in *dPIP5K^18^*,dPIP5K^L-K189A^, which is catalytically inactive (Fig S2B)was unable to rescue the ERG phenotype (Fig 2F). These findings demonstrate that dPIP5K^L^ is sufficient to sustain a normal electrical response to light, a function that is dependent on its kinase activity.

### dPIP5K^L^ is essential for normal phototransduction

Next, we checked whether dPIP5K^L^ is essential for phototransduction. To answer this question, we designed short hairpin constructs that would specifically target and knockdown dPIP5K^L^ by targeting the exon coding for the unique C terminus region of dPIP5K^L^**[20,21,22]**. To test the effectiveness of these hairpin constructs a Western blot was performed from *Drosophila* S2R+ cells co-transfected with *dPIP5K^L^* and each of the hairpin constructs to check for dPIP5K^L^ protein knockdown. Of the three hairpins we designed, construct 1 showed the highest level of knockdown (Fig S3A) and hence we generated transgenic fly lines with it. We used a number of different eye specific enhancers that differ in terms of strength and timing of expression [GMR-3^rd^ larval instar onwards and Rh-70% pupal development onwards] to deplete *dPIP5K^L^* transcript (Fig S3B) and this protein isoform was also depleted (Fig 3A). The transcript level of *dPIP5K^S^* was not affected (Fig S3C) We observed that with increasing degree of dPIP5K^L^ depletion, the amplitude of the ERG response diminishes (Fig 3B,C). Further, upon knockdown with *Rh1* enhancer, which starts expressing from 70 % pupal development, about 70 percent of the eclosed flies, showed a lower ERG amplitude when compared to controls but with normal on/off transients (that report normal synaptic transmission), implying that the diminished amplitude upon depletion of dPIP5K^L^ is not a downstream consequence of altered synaptic transmission but rather, the protein is required to sustain a normal receptor potential during the light response. These experiments demonstrate that dPIP5K^L^ is both necessary and sufficient to sustain a normal electrical response to light.

**Figure 3:**
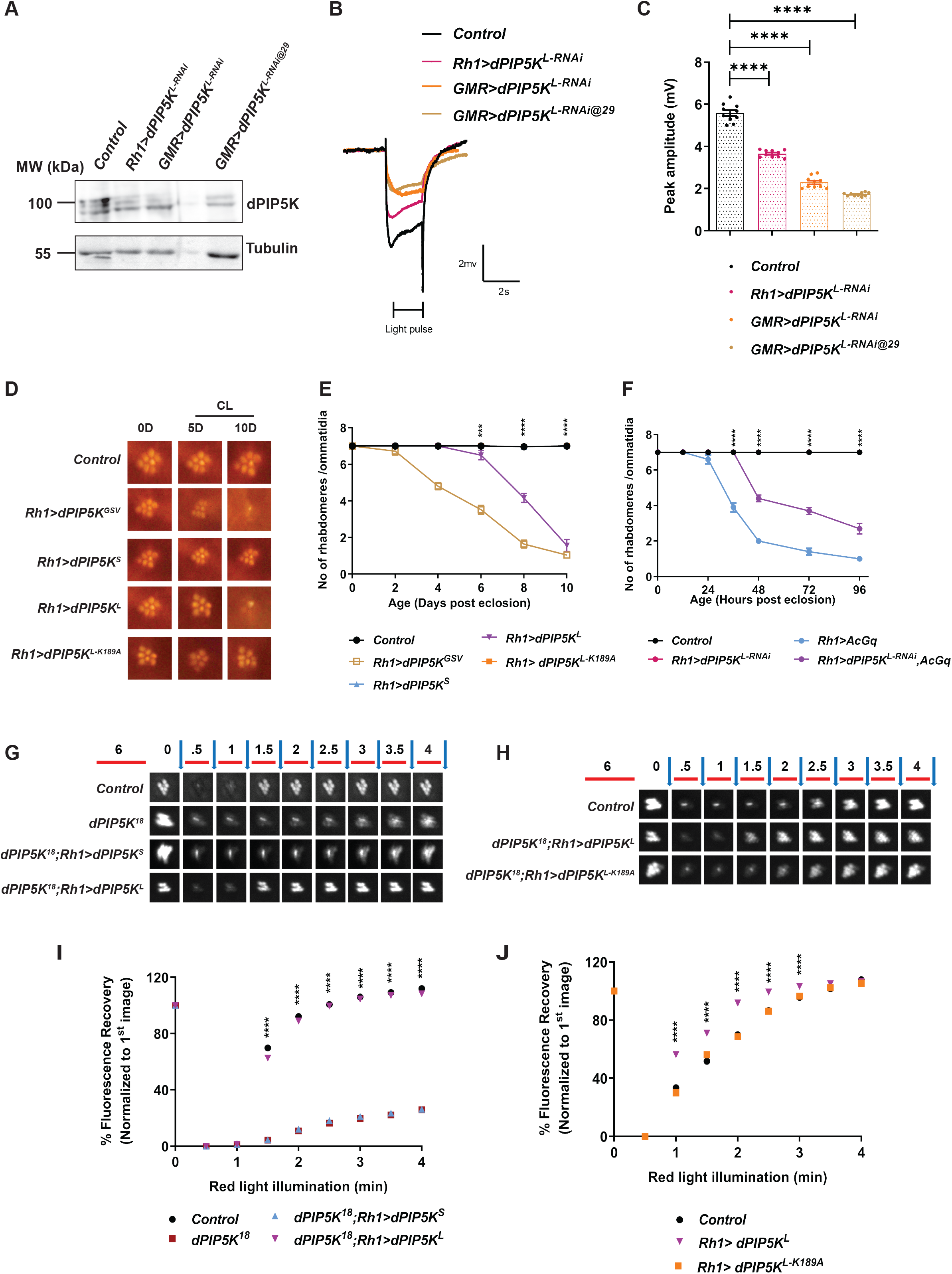
The kinase activity of dPIP5K^L^ is essential for phototransduction. **(A)** Western blot from 1- day old fly retinae showing the degree of knockdown of dPIP5K^L^ (100kDA band) in the mentioned genotypes upon expression of an RNAi to this isoform; blot was probed with dPIP5K antibody. Tubulin at 55kDa is used as loading control. **(B)** Representative ERG trace from 1-day old flies of the indicated genotypes. The duration of the stimulus-a 2s flash of green light is shown Y axis shows response amplitude in mV, X axis shows duration of the recording (s). **(D)** Quantification of the peak amplitude for ERG response in each of the mentioned genotypes.Y axis represents peak amplitude and X axis shows the genotypes. Each data point represents an individual fly tested (n=10 flies). Error bars represent S.E.M. **(D)** Representative optical neutralization images for the mentioned genotypes. Flies were grown in constant light at 2500 lux; age in days (D) is mentioned at the top of each panel. 0D is the day of eclosion, 5D-5 days and 10D-10 days post eclosion, CL-constant light **(E,F)** Graph showing quantification for the time course of retinal degeneration on overexpression of the mentioned transgenes under constant light conditions. (Light intensity:[E] 2500 lux; [F] 5500 lux).10 ommatidia from 5 separate flies of each genotype were scored and plotted. Y-axis is the number of intact rhabdomeres/ommatidium. The maximum value possible is 7. X-axis is the age of the flies post-eclosion. Error bars represent S.E.M. **(G, H)** Deep pseudopupil imaging of PIP_2_ levels in the microvillar membrane of photoreceptors. The fluorescence of the PH-PLCδ∷GFP probe is depicted. The protocol used is represented: Red light illumination periods are shown as red bars, with the numbers on top representing illumination time in minutes. Flashes of blue light (blue arrows) used for image capture are also depicted. Representative fluorescent image captured for each time point for the mentioned genotypes is shown. **(I, J)** Graph showing quantification of recovery kinetics of the fluorescent deep pseudopupil with time in flies expressing the PI(4,5)P_2_ reporter PH-PLCδ::GFP. Y-axis represents the mean fluorescence intensity of the pseudo pupil image at each time point expressed as percentage of the intensity of first image acquired, X-axis represents the red light illumination period in minutes (n = 10 flies). Error bars represent S.E.M. Data information: XY plots and scatter dot plots with mean±S.E.M are shown. Statistical tests: (C) One way Anova with post hoc Tukey’s multiple pairwise comparison. (E,F,I,J) Two-way ANOVA grouped analysis with Bonferroni’s post multiple comparison tests to compare means. ***p<0.001; ****p<0.0001. The statistical significance for the following data pairs is represented on the graphs: (E) *Control vs Rh1>dPIP5K^L^;* (F) *Rh1>AcG_q_ vs Rh1>dPIP5K^L^, AcG_q_*; (I) *dPIP5K^18^ vs dPIP5K^18^;Rh1>dPIP5K^L^*; (J) *Control vs Rh1>dPIP5K^L^*

Light dependent retinal degeneration is a phenotype frequently associated with changes in gene products that are a part of phototransduction [reviewed in (Raghu et al., 2012)]. When all isoforms of dPIP5K (*Rh1>dPIP5K^GSV^*) are overexpressed in photoreceptors using *dPIP5K^GSV^* (Toba et al., 1999), a strong light dependent retinal degeneration is seen (Fig 3D,E). To test the contribution of each isoform to this phenotype, in this study, we overexpressed them individually and found that overexpression of dPIP5K^L^ (*Rh1>dPIP5K^L^*) led to light dependent retinal degeneration (Fig 3D, E), whereas overexpression of dPIP5K^S^ did not result in this phenotype (*Rh1>dPIP5K^S^*) (Fig 3 D, E). The ability of dPIP5K^L^ overexpression to trigger retinal degeneration was dependent on kinase activity of the enzyme since overexpression of dPIP5K^L-K189A^, a kinase dead version did not show any degeneration (Fig 3 D,E).

The heterotrimeric G-protein α subunit G_q_, is essential for PLCβ activity which in turn is required for phototransduction (Scott et al., 1995) and expression of a constitutively active form of G_q_ (AcG_q_) activates phototransduction and accelerates retinal degeneration (Lee et al., 1994). If dPIP5K^L^ generates the PI(4,5)P_2_ pool required for PLCβ activity, then one might predict that depletion of this isoform might suppress the effects of AcG_q_ expression. When AcG_q_ (Ratnaparkhi et al., 2002) was expressed in photoreceptors (*Rh1>AcG_q_*),we found that when reared in light, these photoreceptors underwent retinal degeneration (Fig 3F); this retinal degeneration was slowed down when dPIP5K^L^ was depleted in photoreceptors (Fig 3F).

### dPIP5K^L^ is necessary and sufficient to support PI(4,5)P_2_ turnover during phototransduction

dPIP5K^L^ and dPIP5K^S^ showed a differential ability to rescue the electrical response to light, a process that depends critically on normal PI(4,5)P_2_ turnover during phototransduction. It has previously been reported that in the loss-of-function mutant of *dPIP5K* (*dPIP5K^18^*), the kinetics of PI(4,5)P_2_ turnover is severely impaired (Chakrabarti et al., 2015). To test if this differential rescue of the ERG amplitude by dPIP5K^L^ and dPIP5K^S^ correlated with their ability to support PIP_2_ turnover, we measured PIP_2_ levels at the microvillar membrane *in vivo*. When *dPIP5K^L^* was selectively reconstituted in *dPIP5K^18^*, the delayed kinetics of PI(4,5)P_2_ re-synthesis in this mutant was rescued; by contrast selective reconstitution of *dPIP5K^18^* with *dPIP5K^S^* failed to do so (Fig 3G,I). Moreover, upon overexpression of dPIP5K^L^ in otherwise wild type flies, the kinetics of PI(4,5)P_2_ re-synthesis was accelerated; this was dependent on the kinase activity of dPIP5K^L^ since overexpression of dPIP5K^L-K189A^(*Rh1> dPIP5K^L-K189A^*) did not accelerate the resynthesis of PI(4,5)P_2_ levels (Fig 3H, J). These observations indicate that dPIP5K^L^ is both necessary and sufficient to control PIP_2_ resynthesis following light induced PLCβ activity.

### Septin 7 depletion alters the light response and PI(4,5)P_2_ turnover

Septin monomers have been shown to bind PI(4,5)P_2_ (Zhang et al., 1999) and this binding can modulate the activity of this class of proteins (Bertin et al., 2010; Tanaka-Takiguchi et al., 2009) raising the attractive possibility that they may form the molecular link between PLCβ activity and PI(4,5)P_2_ levels at the plasma membrane. We studied the class 7 septins, which in *Drosophila* are encoded by a single type of monomer encoded by the peanut (*pnut*) gene (Neufeld and Rubin, 1994); we tested the effect of depleting *pnut* on phototransduction. For this we used *pnut^XP^*, a null allele of this gene; since *pnut^XP^* flies are not homozygous adult viable, we used *pnut^XP^*/+ for our experiments. In Western blots from *Drosophila* heads, *pnut^XP^*/+ flies show ca. 50% of the protein levels as compared to wild type controls (Fig 4A). We found that with equal intensities of stimulation, *pnut^XP^*/+ flies have an enhanced ERG response compared to controls (Fig 4B) and the sensitivity of these flies to light was substantially enhanced relative to controls (Fig 4C). These findings were re-capitulated by depletion of *pnut* in photoreceptors using RNAi (*Rh1>pnut^RNAi^*) (Fig 4A); that also resulted in enhanced sensitivity to light (Fig 4D). We also measured PI(4,5)P_2_ turnover in *pnut* depleted photoreceptors; resting levels of PI(4,5)P_2_ were no different between *pnut^XP^*/+ and controls (Fig 4E); however, following PLCβ activation by illumination, PI(4,5)P_2_ levels recovered much faster in *pnut^XP^*/+ compared to controls (Fig 4F). Thus, loss of septin7 activity enhances PI(4,5)P_2_ turnover during PLCβ signalling in photoreceptors.

**Figure 4:**
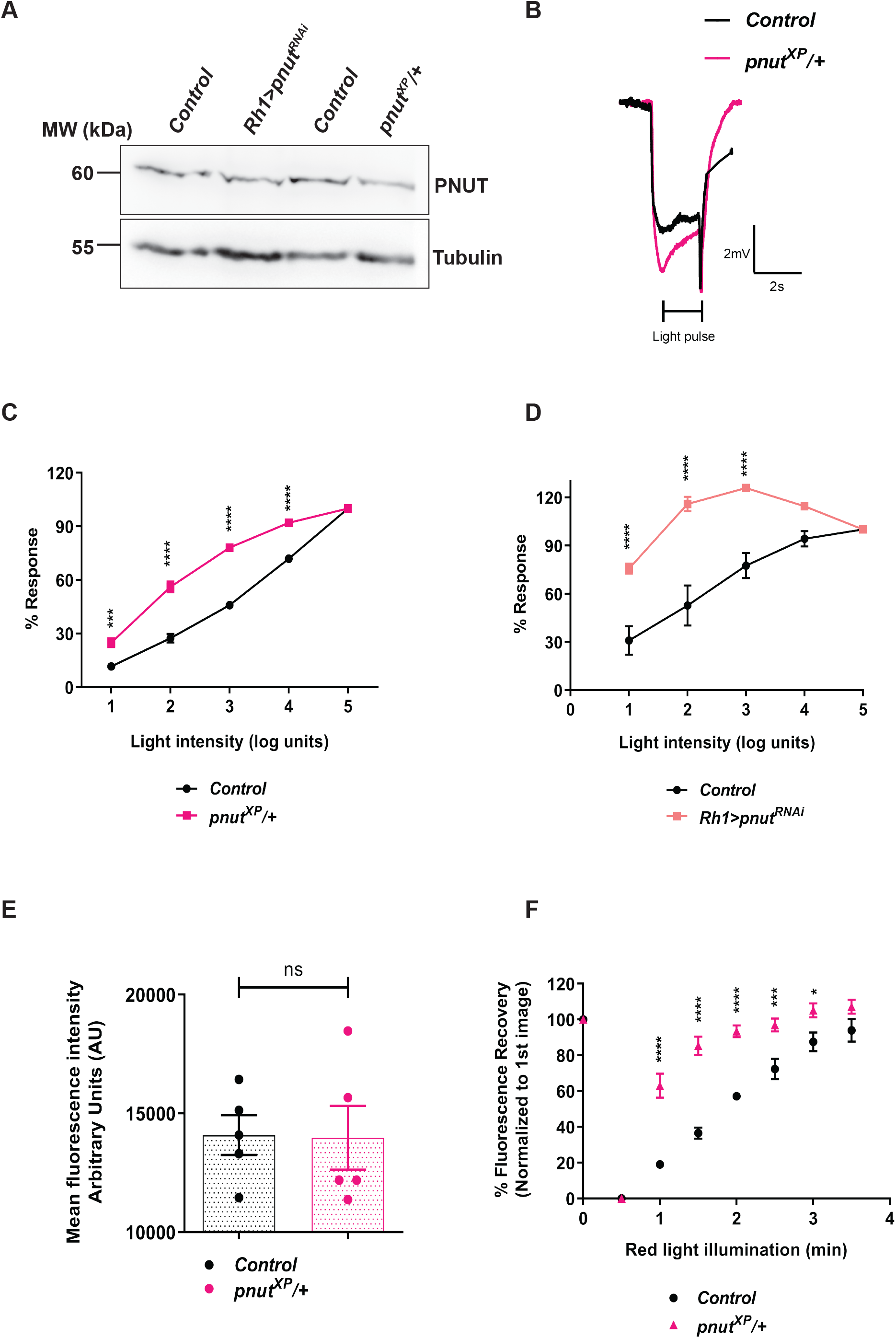
*pnut* depletion alters the light response and PIP_2_ turnover. **(A)** Western blot from retinal extracts of 1- day old flies of the mentioned genotypes. Anti-PNUT detects a band at 60kDa. Tubulin at 55kDa is used as loading control. **(B)** Representative ERG trace from 1- day old flies of the indicated genotypes. The 2s stimulus with a flash of green light is shown. Y axis shows response amplitude (mv), X axis indicates duration of the recording (s). **(C, D)** Light intensity response comparison between: **(C)** *Control* and *pnut^XP/+^* flies **(D)***Control* and *Rh1>pnut^RNAi^* xsxsflies. The X-axis represents increasing light intensity in log units, Y-axis is the peak response amplitude at each intensity normalized to the response at the maximum intensity (n=5). Error bars represent S.E.M. **(E)** Quantification of the mean fluorescence intensity of the deep pseudopupil image in flies expressing the PI(4,5)P_2_ reporter PH-PLCδ∷GFP. Y-axis represents mean fluorescence intensity represented in arbitrary units, X-axis shows the genotypes (n=5). Error bars represent S.E.M. **(F)** Graph showing quantification of recovery kinetics of fluorescent pseudopupil with time in flies expressing the PI(4,5)P_2_ reporter PH-PLCδ∷GFP. Y-axis represents the mean fluorescence intensity of the pseudopupil expressed as percentage of the intensity of first image acquired, X-axis represents the red light illumination period in minutes (n = 5 flies). Error bars represent S.E.M. Data information: XY plots and scatter dot plots with mean±S.E.M are shown. Statistical tests: (E) two tailed unpaired t test with Welch correction. (C,D,F) Two-way ANOVA grouped analysis with Bonferroni’s post multiple comparison tests to compare means.ns-Not significant; *p<0.05; ***p<0.001; ****p<0.0001.

### Septin Interacting Protein 1(Sip1) is an activator of dPIP5K^L^

Given the specific role of dPIP5K^L^ in PI(4,5)P_2_ synthesis during PLCβ signalling, it is interesting to consider how the activity of the enzyme might be regulated during light-induced PLCβ mediated signalling. To identify molecules that might regulate dPIP5K^L^ activity, we considered proteins that are shown to interact with septins. One of these was Septin Interacting Protein 1 (Sip1), first identified in an interaction screen for PNUT/Septin 7 interacting proteins (Shih et al., 2002). Septins are proteins localized to the plasma membrane where several members have been shown to bind PI(4,5)P_2_ (Zhang et al., 1999)and some members have been implicated in receptor activated, PLCβ mediated calcium signalling at this location (Deb et al., 2016; Sharma et al., 2013). Therefore, these molecules may offer a conceptual link between PLCβ mediated PI(4,5)P_2_ hydrolysis at the plasma membrane and dPIP5K^L^ mediated resynthesis of this lipid. If Sip1 is an activator of dPIP5K^L^, depletion of Sip1 in photoreceptors might be expected to phenocopy depletion of dPIP5K^L^. We measured the light response of flies where Sip1 had been depleted in photoreceptors using RNAi (*Rh1>sip1^RNAi^*); this resulted in a reduction of the light response, phenocopying that seen with dPIP5K^L^ knockdown alone (Fig 5A). Importantly, on simultaneous depletion of both Sip1 and dPIP5K^L^, the reduction in ERG amplitude was no greater than the reduction seen when dPIP5K^L^ alone was depleted (Fig 5A). Overexpression of dPIP5K^L^results in a light-dependent retinal degeneration that requires the catalytic activity of the enzyme (Fig 3 D,E). To test if Sip1 was required for dPIP5K^L^ activity, we depleted Sip1 in flies overexpressing dPIP5K^L^. Under these conditions, Sip1 depletion rescued the retinal degeneration induced by dPIP5K^L^over expression (Fig 5 B,C). Lastly, we tested if Sip1 depletion alters the rate of PI(4,5)P_2_ turnover during PLCβ stimulation and found that knockdown of Sip1 significantly slows the kinetics of PI(4,5)P_2_ resynthesis (Fig 5D). Together, these observations support the model that Sip1 can activate dPIP5K^L^ activity *in vivo*.

**Figure 5:**
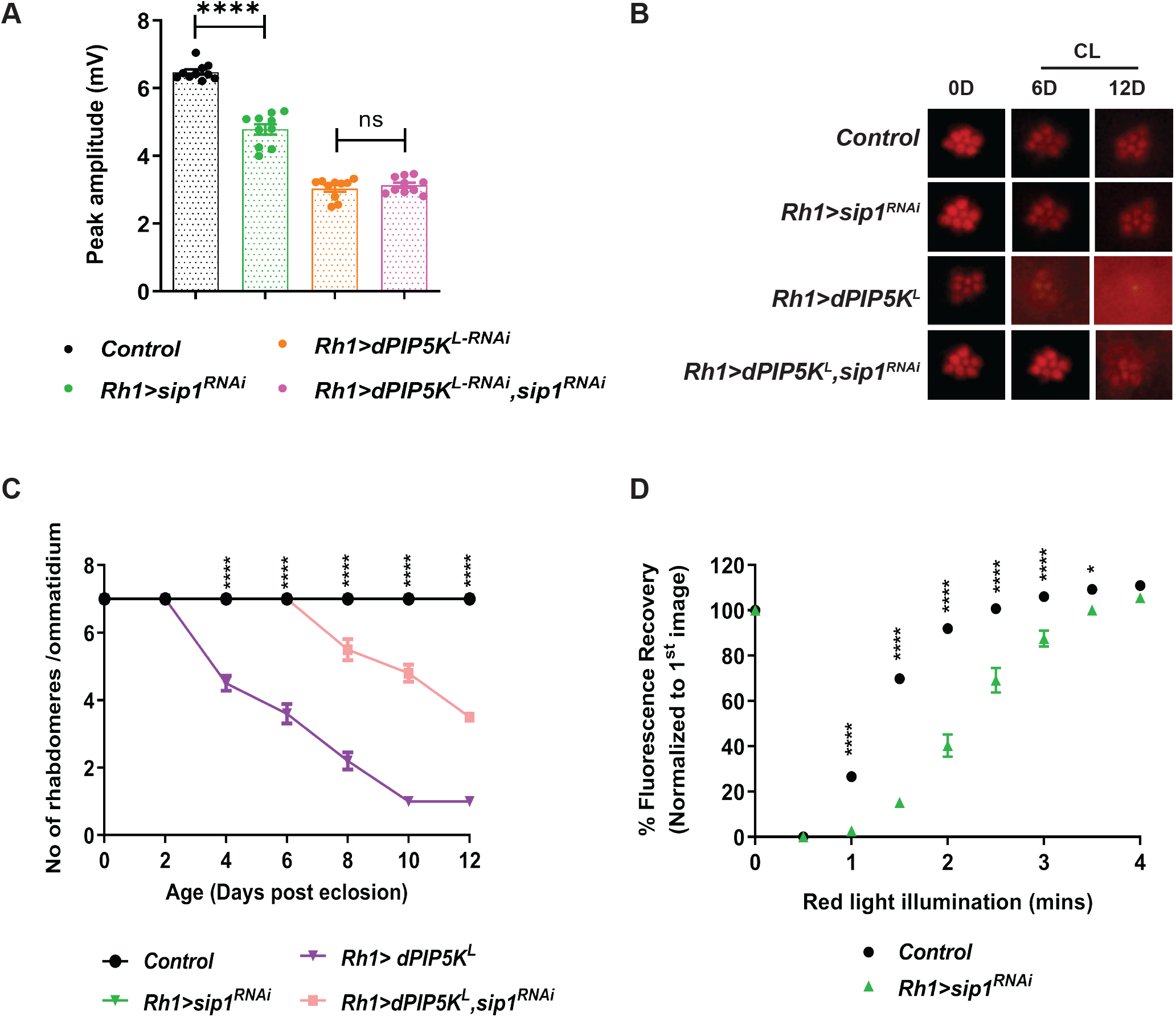
Sip1 acts as a regulator of dPIP5K^L^. **(A)** Quantification of the peak amplitude for ERG response in each of the mentioned genotypes. Y axis represents peak amplitude and X axis shows the genotypes. Each data point represents an individual fly tested. (n=10 flies). Error bars represent S.E.M. **(B)** Representative optical neutralization images for the mentioned genotypes. Flies were grown in constant light at 2500 lux; age in days (D) is mentioned at the top of each panel. 0D is the day of eclosion, 6D-6 days and 12D-12 days post eclosion, CL-constant light. **(C)** Graph showing quantification for the time course of retinal degeneration on overexpression of the mentioned transgenes under constant light at 2500 lux. 10 ommatidia from 5 separate flies of each genotype were scored and plotted. Y-axis is the number of intact rhabdomeres/ommatidium. The maximum value possible is 7. X-axis is the age of the flies post-eclosion. Error bars represent S.E.M. **(D)** Graph showing quantification of recovery kinetics of the fluorescent deep pseudopupil with time in flies expressing the PI(4,5)P_2_ reporter PH-PLCδ∷GFP. Y-axis represents the mean fluorescence intensity of the pseudo pupil image at each time point expressed as percentage of the intensity of first image acquired, X-axis represents the red light illumination period in minutes (n = 10 flies). Error bars represent S.E.M. Data information: XY plots and scatter dot plots with mean±S.E.M are shown. Statistical tests: (A) One way Anova with post hoc Tukey’s multiple pairwise comparison (C,D) Two-way ANOVA grouped analysis with Bonferroni’s post multiple comparison tests to compare means. ns-Not significant; *p<0.05; ****p<0.0001. The statistical significance for the following data pairs is represented on graph (C) *Rh1>dPIP5K^L^* vs *Rh1dPIP5K^L^,sip1^RNAi^*

### Septin 7 depletion can suppress the effects of both dPIP5K^L^ and Sip1

We tested if *pnut* depletion can impact the consequences of Sip1 depletion. When Sip1 depletion was performed in photoreceptors that were also *pnut^XP^*/+, the reduced ERG amplitude of Sip1 depletion was partially rescued (Fig 6A). Likewise, when dPIP5K^L^ was depleted in *pnut^XP^*/+ flies, the reduction in ERG amplitude normally seen when dPIP5K^L^ is depleted was partially rescued (Fig 6B). To test the relationship of this genetic interaction seen at the level of the ERG amplitude to PI(4,5)P_2_ turnover, we measured PI(4,5)P_2_ turnover kinetics comparing *pnut^XP^/+* with dPIP5K^L^ knockdown. We found that that depletion of *pnut* reversed the delayed PI(4,5)P_2_ recovery kinetics seen in flies with knock down of dPIP5K^L^ alone(Fig 6C). If PNUT and dPIP5K^L^ regulate the turnover of PI(4,5)P_2_ during PLCβ signalling, they could be expected to co-localize at or close to the plasma membrane. We studied the localization of these protein in *Drosophila* S2R^+^ cells; in these cells, endogenous PNUT was found at the plasma membrane and when transfected into these cells, dPIP5K^L^ was also found at this location (Fig 6D). Since we had strong evidence for a negative interaction between PNUT and dPIP5K^L^, to directly check for this we did an *in vitro* kinase assay from *Drosophila* S2R+ cells expressing dPIP5K^L^, where PNUT had been depleted using dsRNA against *pnut*. Under these conditions, the activity of dPIP5K^L^ was significantly increased as compared to control cells (treated with dsRNA against GFP) (Fig 6 E,F).

**Figure 6:**
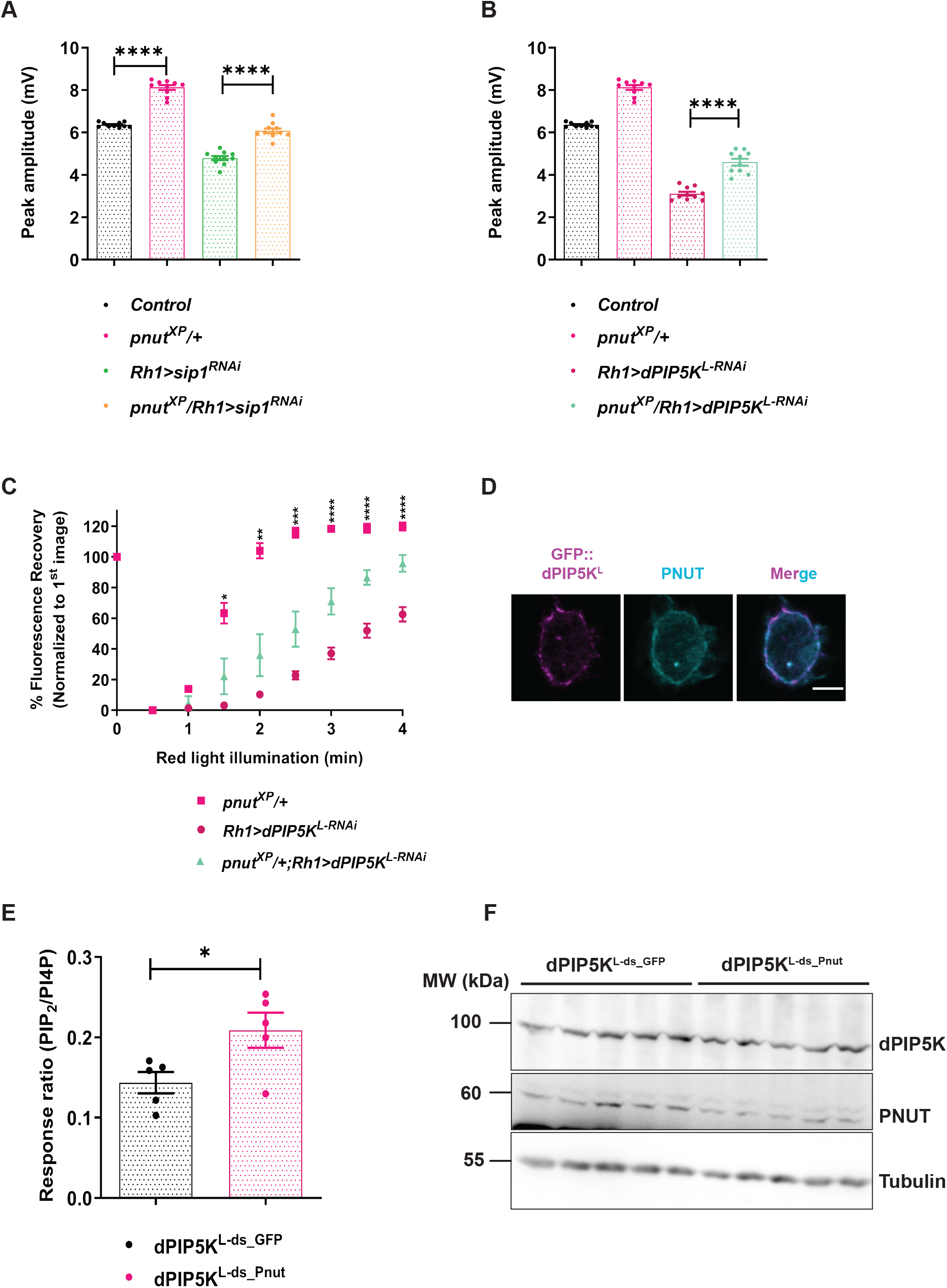
*pnut* depletion can suppress the effects of both dPIP5K^L^ and Sip1. **(A, B)** Quantification of the peak amplitude for ERG response in each of the mentioned genotypes. A and B represent data from the same experiment with common *Control* and *pnut^XP^/+* animals. The data acquired has been represented as two graphs for ease of comparison. Y axis represents peak amplitude and X axis shows the genotypes. Each data point represents an individual fly tested (n=10 flies). Error bars represent S.E.M. **(C)** Graph quantifying the recovery kinetics of fluorescent pseudo pupil with time in flies expressing the PI(4,5)P_2_ reporter PH-PLCδ ∷GFP: Y-axis represents the mean fluorescence intensity of the pseudo pupil image expressed as percentage of the intensity of first image acquired, X-axis represents the red light illumination period in minutes (n = 10 flies). Error bars represent S.E.M. **(D)** Representative confocal images of S2R+ cells transfected with *GFP:: PIP5K^L^*. Localization of endogenous PNUT is detected using the anti-PNUT antibody; GFP∷PIP5K^L^ is detected using the anti GFP antibody. Scale bar: 5μm. **(E)** Response ratio of product:substrate [37:4-PIP_2_:37:4-PI4P] in an *in vitro* kinase assay with cell free lysate prepared from S2R+ cells transfected with dPIP5K^L^, and treated either with control dsRNA against GFP (dPIP5K^L-ds_GFP^), or with dsRNA against PNUT (dPIP5K^L-ds_Pnut^) (n=5 biological replicates). This experiment was repeated twice. Error bars represent S.E.M. **(F)** Western blots showing expression of PNUT from S2R+ cell extract transfected with *dPIP5K^L^*, and treated with dsRNA either against PNUT or control GFP. PNUT is detected at 60 kDa. Knockdown of PNUT can be seen in ds_PNUT treated cells (dPIP5K^L-ds_Pnut^) as compared to control ds_GFP treated cells(dPIP5K^L-ds_GFP^). dPIP5K^L^ (at 100kDa) is expressed equivalently in both cases. Tubulin at 55kDa is used as loading control. Data information: XY plots and scatter dot plots with mean±S.E.M are shown. Statistical tests: (E) two tailed unpaired t test with Welch correction. (A,B) One way Anova with post hoc Tukey’s multiple pairwise comparison. (C) Two-way ANOVA grouped analysis with Bonferroni’s post multiple comparison tests to compare means. *pμ0.05; **pμ0.01; ***pμ0.001; ****pμ0.0001. The statistical significance for the following data pairs is represented on graph (C) *Rh1>dPIP5K^L-RNAi^ vs pnut^XP^/+; Rh1 dPIP5K^L-RNAi^*

**Figure 7:**
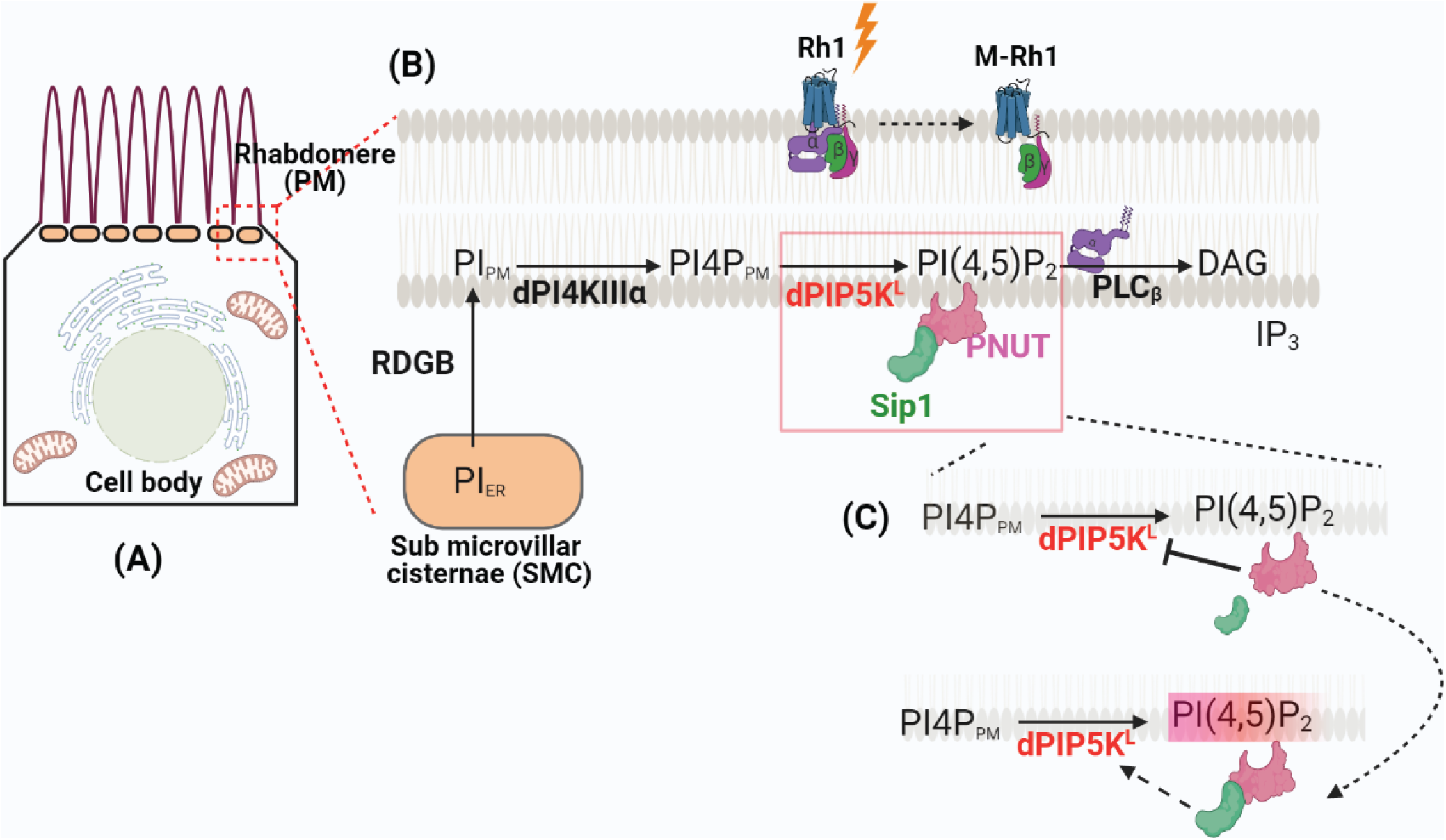
Model for PNUT/Sip1 mediated regulation of PI(4,5)P_2_ synthesis during G-protein coupled PLC signaling in *Drosophila* photoreceptors. (**A**) A cartoon showing cross section of a single *Drosophila* photoreceptor cell with rhabdomere (the apical plasma membrane, **PM**), sub-microvillar cisternae (SMC) which is a modified smooth ER compartment and cell body. **(B)**The area indicated by the red box is expanded to show the biochemical reactions involved in the resynthesis of PI(4,5)P_2_. Protein and lipid components are depicted according to the compartment in which they are distributed. During phototransduction, upon light stimulation, activated PLC (PLCβ) hydrolyses the PI(4,5)P_2_ produced by dPIP5K^L^, depleting the PI(4,5)P_2_ at the plasma membrane. **(C)**PNUT acts as a negative regulator of dPIP5K^L^ activity, thus regulating the levels of PI(4,5)P_2_ present at the microvillar PM. The change in PI(4,5)P_2_ levels at the PM are sensed by the PNUT molecules. Sip1 (septin interacting protein) physically interact with PNUT to transmit this information to dPIP5K^L^, resulting in the activation of dPIP5K^L^ to maintain PI(4,5)P_2_ hemostasis at the photoreceptor membrane. This illustration was created with BioRender.com

## Discussion

The activation of PLC is a widespread mechanism of signalling triggered when plasma membrane receptors bind their cognate ligands. Given that the substrate of PLC, PI(4,5)P_2_ is present at the plasma membrane in low amounts, PLC activation can rapidly deplete this lipid leading to loss of signalling function in cells. This situation may arise in settings such as neurons where GPCRs activate PLC at high rates or T-cell receptor activation where sustained activation of PLC occurs over a long period of time. In order to avoid the depletion of PI(4,5)P_2_, leading to loss of signalling, it is essential that the resynthesis of this lipid is coupled with its consumption by PLC activity. To achieve this, conceptually, cells require a mechanism to sense the drop in PI(4,5)P_2_ levels and convey the requirement for resynthesis to the enzymatic machinery for synthesis whose activity can then restore PI(4,5)P_2_ levels at the plasma membrane.

In eukaryotic cells, the synthesis of PI(4,5)P_2_ is primarily mediated by the PIP5K class of enzymes [reviewed in (Kolay et al., 2016)] and in some cells pools of PI(4,5)P_2_ specifically used during PLC signalling are synthesized by specific isoforms of PIP5K (Chakrabarti et al., 2015). How do such isoforms sense the drop in PI(4,5)P_2_ and trigger their activity to resynthesize this lipid ? In this study we report a splice variant of dPIP5K (dPIP5K^L^), that is both necessary and sufficient to support PI(4,5)P_2_ turnover during *Drosophila* phototransduction, a process in which the plasma membrane of photoreceptors experience high rates of PLC activation. Although the dPIP5K gene is alternately spliced to give two isoforms dPIP5K^S^ and dPIP5K^L^, only the latter is enzymatically active *in vitro* and is able to support both PLC signaling *in vivo*. Consistent with its function during phototransduction, we find that dPIP5K^L^expression is enriched in *Drosophila* photoreceptors and the protein localizes to the microvillar membrane, the sub-cellular site of phototransduction, where most of the proteins involved in this signalling process are also localized. A number of findings reported here (Fig 2, 3) strongly indicate that the kinase activity of dPIP5K^L^ is required for its ability to support PLC signaling *in vivo* and is well correlated with the ability of this polypeptide to synthesize PI(4,5)P_2_ both *in vitro* and *in vivo*. By contrast, dPIP5K^S^ did not show kinase activity *in vitro* and was not able to support PLC signalling in *Drosophila* photoreceptors *in vivo*. Given that the principal difference between dPIP5K^L^ and dPIP5K^S^ is the C-terminal extension of the longer isoform, it would seem reasonable that this unique C-terminal region of 273 amino acids will be important in the control of enzyme activity and ability to support function *in vivo*. A conceptually similar observation has been reported in the case of a mammalian PIP5Kγ isoform, where the binding of FERM domain of talin to the C-terminal region of the enzyme has been reported to link this isoform to ongoing integrin signalling (Di Paolo et al., 2002)(Ling et al., 2002).

In order for dPIP5K^L^ to be activated during ongoing PLC signalling, there is a requirement for a molecule that can detect plasma membrane levels of PI(4,5)P_2_ and then convey a drop in its levels to the PIP5K. Septins are once such class of molecules that localize to the plasma membrane and bind PI(4,5)P_2_ and could in principle act as a sensor of PI(4,5)P_2_ levels. Using a mutant in *pnut*, the only septin 7 ortholog in *Drosophila*, we found that depletion of this molecule results in an enhanced light response and accelerates the recovery of PI(4,5)P_2_ levels at the microvillar plasma membrane following PLC activation. This observation implies that septin function is important for the regulation of plasma membrane PI(4,5)P_2_ levels following PLC activation at the plasma membrane. Altered PI(4,5)P_2_ distribution has been observed at the plasma membrane in mammalian cells in the context of G_q_-PLC mediated calcium signalling following depletion of specific septin family members although the mechanism underlying this has not been discovered (Sharma et al., 2013). We observed that the accelerated PI(4,5)P_2_ resynthesis seen in *pnut^XP/+^*could be reverted back towards wild type by depletion of dPIP5K^L^ (Fig 6C) further strengthening the idea that septins play a role in PI(4,5)P_2_ resynthesis during PLC signalling. We observed that in *Drosophila* S2R+ cells both PNUT and dPIP5K^L^ are localized to the plasma membrane (Fig 6D), consistent with a role for septin dependent regulation of PIP5K function. These findings strongly support a role for septins as a regulator of PI(4,5)P_2_ turnover during PLC signalling.

If septins regulate dPIP5K^L^ activity, what is the mechanism by which they do so ? Although PNUT and dPIP5K^L^ both localize to the plasma membrane, what is the mechanism by which PNUT regulates the activity of dPIP5K^L^? Although a direct physical interaction between these two molecules is a possible mechanism, evidence for this is presently not available. One possible mechanism is an intermediate molecule that is known to interact with septins and may convey a signal to activate dPIP5K^L^. Several septin interacting proteins have been identified (Shih et al., 2002) and molecules such as these could in principle serve to transduce PI(4,5)P_2_ sensing by septins into dPIP5K^L^ activation; a prediction is that for any such transducer, it’s depletion might phenocopy that of dPIP5K^L^ depletion. In this study, we found that depletion of Sip1, a septin interacting protein (Shih et al., 2002) phenocopied multiples aspects of dPIP5K^L^ depletion (Fig 5,6) including: (i)reduced electrical response to light (ii) slower rate of PI(4,5)P_2_ resynthesis. In addition, Sip1 depletion was able to suppress the retinal degeneration phenotype seen when dPIP5K^L^ is over-expressed. Lastly Sip1 depletion was able to reverse the phenotype of *punt^XP/+^* in a manner similar to the ability of dPIP5K^L^ to suppress the effects of *pnut* depletion. Together these findings make a compelling case for Sip1 acting to transduce signals from PNUT to dPIP5K^L^. These findings suggest a general mechanism by which PI(4,5)P_2_ levels at the plasma membrane might be sustained during receptor activated PLC signalling.

## Acknowledgements

This work was supported by the Department of Atomic Energy, Government of India, under Project Identification No. RTI 4006, a Wellcome-DBT India Alliance Senior Fellowship (IA/S/14/2/501540) to PR. We thank the transgenic fly facility, Central Imaging Facility and mass spectrometry Facility at NCBS for support. Sourav Kolay was supported by a fellowship from the Council for Scientific and Industrial Research, Government of India.

## Materials and Methods

### Fly stocks

Fly stocks were maintained in 25^0^C laboratory incubators with 50% relative humidity and no internal illumination. All flies were raised on standard corn meal media containing 1.5% yeast. For all the constant light experiments, flies were grown in an incubator with constant illumination from a white light source for the required time periods.

The wildtype used was Red Oregon-R (ROR). Gal4 UAS system was used for targeted expression of transgenic constructs. The following fly alleles and insertions obtained for the experiments are described here: so^D^ (Bloomington Stock #4287), dPIP5K over expression line GS200386 (DGRC-Kyoto), dPIP5K^18^ (Chakrabarti et al., 2015), dPIP5K^L^, dPIP5K^L-K189A^, dPIP5K^L^-RNAi,dPIP5K^S^+HA and dPIP5K^L^∷GFP (generated during this study), *pnut^XP^/+*, UAS-AcG_q_ (Gaiti Hasan, NCBS Bangalore), Sip1RNAi (Bloomington stock #56933), PNUTRNAi (VDRC: v11791)

### RNA Extraction and Q-PCR

RNA extraction was done from 1- day old *Drosophila* retinae or heads using TRIzol reagent (Invitrogen). After this, purified RNA was treated with amplification grade DNase I (Invitrogen). Reverse transcription was done using SuperScript II RNase H– Reverse Transcriptase (Invitrogen) and random hexamers (Applied Biosystems). Quantitative PCR (Q-PCR) was performed with the Applied Biosystems 7500 Fast Real Time PCR instrument. Primers were designed at the exon-exon junction following the parameters recommended for Q-PCR. Transcript levels of the ribosomal protein 49 (RP49) were used for normalization across samples. Three separate samples were collected from each genotype, and duplicate measures of each sample were conducted to ensure the consistency of the data. The following primer pairs were used:

RP49 fwd: CGGATCGATATGCTAAGCTGT
RP49 rev: GCGCTTGTTCGATCCGTA
Sktl fwd: CTCATGTCCATGTGTGCGTC
Sktl rev: TTAATGGTGCTCATCAGTG
dPIP5K fwd: AGCAGAGAAAACCGCTTAGG
dPIP5K rev: GGCGATTCACTGACTTATTCC
dPIP5K^L^ fwd: AGAACCGCACACCACTAATCCC
dPIP5K^L^ rev: GCGGTGCATATGGACCATATGG
dPIP5K^S^ fwd: GGACATTGTGAGCGTTTCGG
dPIP5K^S^ rev: TCTGTCCATGTGGGAGTGGA

### Cloning of dPIP5K^L^

Total RNA was isolated from fly retinae using the Trizol reagent, following which the RNA was treated with DNAse I amplification grade for digestion of genomic DNA. Full length cDNA clones were obtained using primer specific to the 3’ end of *dPIP5K^L^* and then cloned into an intermediate vector: pCR-XL-TOPO (TOPO XL PCR Cloning Kit, Invitrogen) and finally subcloned into fly expression vector pUAST-attB using Not1 and Xho1 restriction sites. Transgenic flies were obtained upon microinjection. dPIP5K^L_K189A^ (kinase dead) was obtained by doing a site directed mutagenesis at Lysine (K)189 position in dPIP5K^L^. The following primer pairs were used:

To obtain *dPIP5K^L^* cDNA: TGCCTAGCAATACAATGAGC

To clone dPIP5K^L^ into pUAST-attB:

UAS dPIP5K^L^ Not1_ fwd: GCGGCCGCATGGCCTCCGGCGAT

UAS dPIP5K^L^ Xho1_rev: GAGAGACTCGAGCTAATCCCGTCCCTGTCC

To generate dPIP5K^L_K189A^ kinase dead mutant:

dPIP5K^L_K189A^ fwd: ACGAGTTCATCATAGCGACGGTGCAACACAA
dPIP5K^L_K189A^ rev: ACGAGTTCATCATAGCGACGGTGCAACACAA

### Generation of dPIP5K^L-RNAi^ flies

The Transgenic RNAi Project (TRiP) guidelines were used to synthesize shRNA constructs against dPIP5K^L^, which were then cloned into Walium 20 vector. Transgenic flies were obtained upon microinjection of the constructs. The following primer pair was used:

dPIP5K^L^_RNAi fwd: CTAGCAGTGCGTCCTAGAAGACCTCTACCTAGTTATATTCAAGCATAGGTAGAGGTCTTCTAGGACGCGCG
dPIP5K^L^_RNAi rev: AATTCGCGCGTCCTAGAAGACCTCTACCTATGCTTGAATATAACTAGGTAGAGGTCTTCTAGGACGCACTG

### Electroretinogram

Flies were briefly put on ice to anaesthetize and immobilized at the end of a disposable pipette tip such that the head protruded out. Recordings were done using glass microelectrodes filled with 0.8% w/v NaCl solution. Voltage changes were recorded between electrode placed at the surface of the eye and the ground electrode placed on the thorax. Before recording, flies were dark adapted for 5 minutes. Recordings were done with 2-s flashes of green light stimulus, with 10 stimuli (flashes) per recording and 15s of recovery time between two flashes of light. Green light stimulus was delivered from a LED light source within 5 mm of the fly’s eye through a fibre optic guide. Calibrated neutral density filters were used to vary the intensity of the light source during intensity response experiments. Voltage changes were amplified using a DAM50 amplifier (SYS‐ DAM50, WPI, Florida, USA) and recorded using pCLAMP 10.7(Molecular devices, California, USA). Analysis of traces was performed using Clampfit 10.7. Analysed results were plotted using GraphPad Prism software.

### Pseudopupil assay

To monitor PI (4,5) P_2_ dynamics in live flies, transgenic flies expressing PH-PLCδ∷GFP [PI(4,5)P_2_ biosensor] were anaesthetized and immobilized at the end of a pipette tip using a drop of colorless nail varnish and fixed by clay on the stage of an Olympus IX71 microscope. The fluorescent deep pseudopupil (DPP, a virtual image that sums rhabdomere fluorescence from ~20– 40 adjacent ommatidia) (Chakrabarti et al., 2015; Franceschini et al., 1981) was focused and imaged using a 10X objective. Time-lapse images were taken by exciting GFP using a 90ms flash of blue light and collecting emitted fluorescence. The program used for this purpose was created in Micromanager. Following preparation, flies were adapted in red light for 6 minutes after which the eye was stimulated with a 90 ms flash of blue light (ƛ_max_488nm). The blue light used to excite GFP was also the stimulus to rapidly convert the majority of rhodopsin (R) to metarhodopsin (M) thus activating the phototransduction cascade and triggering depletion of rhabdomeric PI(4,5)P_2_. Between the blue light stimulations, photoreceptors were exposed to long wavelength (red - ƛ_max_660nm) light that reconverts M to R. The recovery in DPP fluorescence intensity with time indicates translocation of the probe from cytoplasm to rhabdomere membrane upon PI(4,5)P_2_ resynthesis. The DPP intensity was measured using ImageJ from NIH (Bethesda, MD, USA). Cross sectional areas of rhabdomeres of R1-R6 photoreceptors were measured and the mean intensity values per unit area were calculated.

### Optical neutralization

Flies of the desired age and reared under the required experimental conditions were decapitated and the heads fixed on a glass slide using a drop of colorless nail polish. Imaging of the photoreceptors was done using a 40X oil immersion objective lens (Olympus BX43). Images were acquired by using CellSens software. For light adaptation, newly eclosed flies were transferred to incubators programmed with constant light settings.

### Scoring retinal degeneration

To calculate a quantitative index of degeneration, 50 ommatidia from 5 different flies of each genotype were counted for each time point. The central photoreceptor, which is UV sensitive, hence did not show any light dependent retinal degeneration, was used as the reference and the rest of the photoreceptors were scored. Each intact rhabdomere was assigned a score of 1, and each degenerated rhabdomere was counted as 0. Hence, the maximum number is 7 and minimum is 1 (representing R7, which is insensitive to white light) Thus, control photoreceptors will have a score of 7 and the genotypes that undergo degeneration will have a score from 1 to 7. Analyzed results were plotted using GraphPad Prism software.

### Western Blotting

Heads or retinae from 1-day-old flies were harvested, and crushed in 2X SDS-PAGE sample buffer followed by boiling at 95°C for 5 minutes. For protein sample preparation from S2R+ cells, 48 hours post transfection, cells were harvested, washed twice with PBS and lysed in 2X SDS-PAGE sample buffer. The genomic DNA was sheared using an insulin syringe followed by boiling the samples at 95°C for 5 minutes. Protein extracts were separated using SDS-PAGE and electro blotted onto nitrocellulose filter membrane [Hybond‐C Extra; (GE Healthcare, Buckinghamshire, UK)], using semidry transfer apparatus (Bio-Rad). Next, the membrane was blocked in 5% Blotto (sc‐2325, Santa Cruz Biotechnology, Texas, USA) in phosphate buffer saline (PBS) with 0.1% Tween 20 (Sigma‐Aldrich) (PBST) for 2 hours at room temperature. Primary antibody incubation was done overnight at 4°C using the appropriate antibody dilutions. The following antibodies were used: anti-α-tubulin (1:4000, DHSB [E7c]), anti dPIP5K (1:1000, lab generated), anti GFP (1:2000, SC [B2]), anti HA (1:1000, CST [6E2]), anti-PNUT (1:250, DSHB [4C9H4-c]). Following this, the membrane was washed thrice in PBST for 10 minutes each and incubated with 1:10000 dilutions of appropriate secondary antibody coupled to horseradish peroxidase (Jackson ImmunoResearch Laboratories) at room temperature for 2 hours. Next, the membrane was washed thrice in PBST for 10 minutes each, developed with ECL reagent (GE Healthcare) and imaged in LAS 400 instrument (GE Healthcare).

### Immunohistochemistry

For Immunohistochemistry, retinae from 1- day old flies were dissected in chilled phosphate buffer saline (PBS), followed by fixation in 4% paraformaldehyde in PBS with 1 mg/ml saponin at room temperature for 30 minutes on orbital shaker. Fixed retinae were washed three times in PBTx (1X PBS + 0.3% TritonX-100) for 10 minutes each. The sample was then blocked using 5% Fetal Bovine Serum in PBTx for 2 hours at room temperature on shaker, after which the sample was incubated with primary antibody in blocking solution overnight at 4°C on a shaker. The following antibodies were used: anti-GFP (1:5000,abcam [ab13970]), anti-HA (1:50, CST [6E2]) Appropriate secondary antibodies conjugated with a fluorophore were used at 1:300 dilutions [Alexa Fluor 488/568/633 IgG, (Molecular Probes)] and incubated for 4 hours at room temperature. Wherever required, during the incubation with secondary antibody, Alexa Fluor 568-phalloidin (1:200, Invitrogen [A12380]) was also added to the tissues to stain the F-actin. After three washes in PBTx, samples were mounted in 70% glycerol in 1X PBS. Whole mounted preparations were imaged on Olympus FV3000 confocal microscope using Plan-Apochromat 60x, NA 1.4 objective (Olympus). Image analysis was performed using ImageJ from NIH (Bethesda, MD, USA).

### Cell Culture, Transfections and dsRNA treatment

*Drosophila* S2R+ cells stably expressing ActGal4 were maintained on Schneider’s media supplemented with 10% non heat inactivated fetal bovine serum(FBS) and contained antibiotics-streptomycin and penicillin (Schneider’s Complete Media, SCM). Cell transfections were carried out using Effectene (301425, Qiagen) as per the manufacturer’s protocol. For *pnut* knockdown experiments, 0.5 million cells were plated in 5 biological replicates each and allowed to settle overnight. Post this, SCM was removed and 300μl of Schneider’s incomplete media (SIM) was added to the cells, followed by addition of 1.875μg of the desired dsRNA and incubated for 1 hour. dsRNA against GFP was used as a control. Post an hour, 300μl of SCM was added. This procedure was repeated post 48 hours, followed by transfections of the cells with dPIP5K^L^ plasmid, as described above. All dsRNA were synthesized using the MEGAscript RNAi kit. (AM1626, Thermo). Primers for dsRNA amplification were synthesized in accordance with the DRSC/TRiP Functional Genomics Resources.

The following primer pairs were used:

dsGFP fwd: TAATACGACTCACTATAGGGATGGTGAGCAAGGGCGAGGAG
dsGFP rev: TAATACGACTCACTATAGGGCTTGTACAGCTCGTCCATGCCG
dsPnut fwd: TAATACGACTCACTATAGGGGAAGGAGTGGGAGGATGTCA
dsPnut rev: TAATACGACTCACTATAGGGTTAGGATCAATTTCGCCTCG

### Lipid kinase assay

#### Cell lysate preparation

S2R+ cells expressing the desired constructs were dislodged from plates, harvested by centrifugation at 1675x***g*** for 5 minutes in a table top centrifuge. Then they were washed twice with PBS, and lysed with lysis buffer (50mM Tris-Cl, 1mM EGTA, 1mM EDTA, 1% Triton X-100, 50mM NaF, 0.27M Sucrose, 0.1% 2-Mercaptoethanol) with freshly added protease inhibitor and phosphatase inhibitor cocktail (Roche). Protein equivalent to 10μg was estimated by Bradford assay.

#### Micelle formation and Kinase assay

600 pmoles of 37:4 PI4P was dried in LoBind 1.5ml Eppendorf tubes along with 20μl 0.5mM Phosphatidylserine (SIGMA) and 200μL of CHCL3 in vacuum for 20 minutes at 949x***g***. After drying of lipids, 50μl of 10mM Tris-HCl (pH 7.4) was added to each tube and sonicated at room temperature for 3 minutes. Next, 50 μl of 2X kinase assay buffer containing 40μM ATP (Roche) was added to the tubes and 10μg equivalent of corresponding lysate was added. Immediately after this, the tubes were transferred to a shaking incubator at 30^0^C for 30 minutes for the kinase reaction to happen. The reaction was stopped by adding 125μl of 2.4N HCl to the reaction tubes. Then 250μl each of CHCl3 and methanol were added. After this, the tubes were vortexed for 2 minutes and centrifuged at 1000x****g**** for 5 minutes to allow a clean phase separation to happen The upper phase was punctured and the lower organic phase was transferred to a fresh LoBind 1.5ml Eppendorf. To this tube 500μl of Lower Phase Wash Solution (LPWS: methanol/1 M hydrochloric acid/chloroform in a ratio of 235/245/15 (vol/vol/vol) was added. It was vortexed for 2 minutes and clean phase separation was obtained by spinning tubes at 1000x***g*** for 5 minutes. Again, the lower organic phase was transferred to a fresh LoBind 1.5ml Eppendorf tube.

#### Derivatization and LC MS/MS

The organic phase obtained after lipid extraction was directly subjected to derivatization using 2M TMS-diazomethane (Acros). 50μl TMS-diazomethane was added to each tube and vortexed gently for 10 minutes at 670x***g***. The reaction was neutralized by 10μl of glacial acetic acid. The samples were dried, and reconstituted in 200μl of methanol. LC MS/MS run method was used as described in (Ghosh et al., 2019). The area under the peaks for individual lipid species was extracted using MultiQuant software. Numerical analysis was done in Microsoft Excel and the graphs were plotted using GraphPad Prism software.

#### Immunofloroscence from S2R+ cells

48 hours post transfection with *dPIP5K^L^∷GFP*, cells were fixed in 2.5% PFA for 20 minutes. Upon fixation, cells were washed thrice with PBTx (1× PBS + 0.3% Triton X-100) for 10 minutes each, permeabilized with 0.37% Igepal (Sigma) in 1× M1 for 13 minutes and then blocked for 1 hour at room temperature in 1× M1 containing 5% FBS and 2 mg/ml BSA. Post blocking, the cells were incubated with primary antibodies in blocking solution overnight at 4°C. The following antibodies were used: anti-GFP (1:5000, Abcam [ab13970]) and anti-PNUT (1:250, DSHB [4C9H4]). Next morning, the cells were washed thoroughly, and incubated with the appropriate secondary antibodies conjugated with fluorophore at 1:300 dilutions [Alexa Fluor 488/568/633 IgG, (Molecular Probes)] and incubated at room temperature for 2 hours. The cells were then washed and imaged on an Olympus FV3000 confocal microscope using Plan-Apochromat 60x, NA 1.4 objective (Olympus). Image analysis was performed using ImageJ from NIH (Bethesda, MD, USA)

#### Statistical Analysis

All statistical analyses were done in GraphPad Prism 8 software. Unpaired two-tailed t-test with Welch correction, One way ANOVA, followed by Tukey’s multiple comparison test or Two-way ANOVA grouped analysis with Bonferroni’s post multiple comparison were carried out where applicable

## Supplementary Figures

**Supplementary Fig S1:**
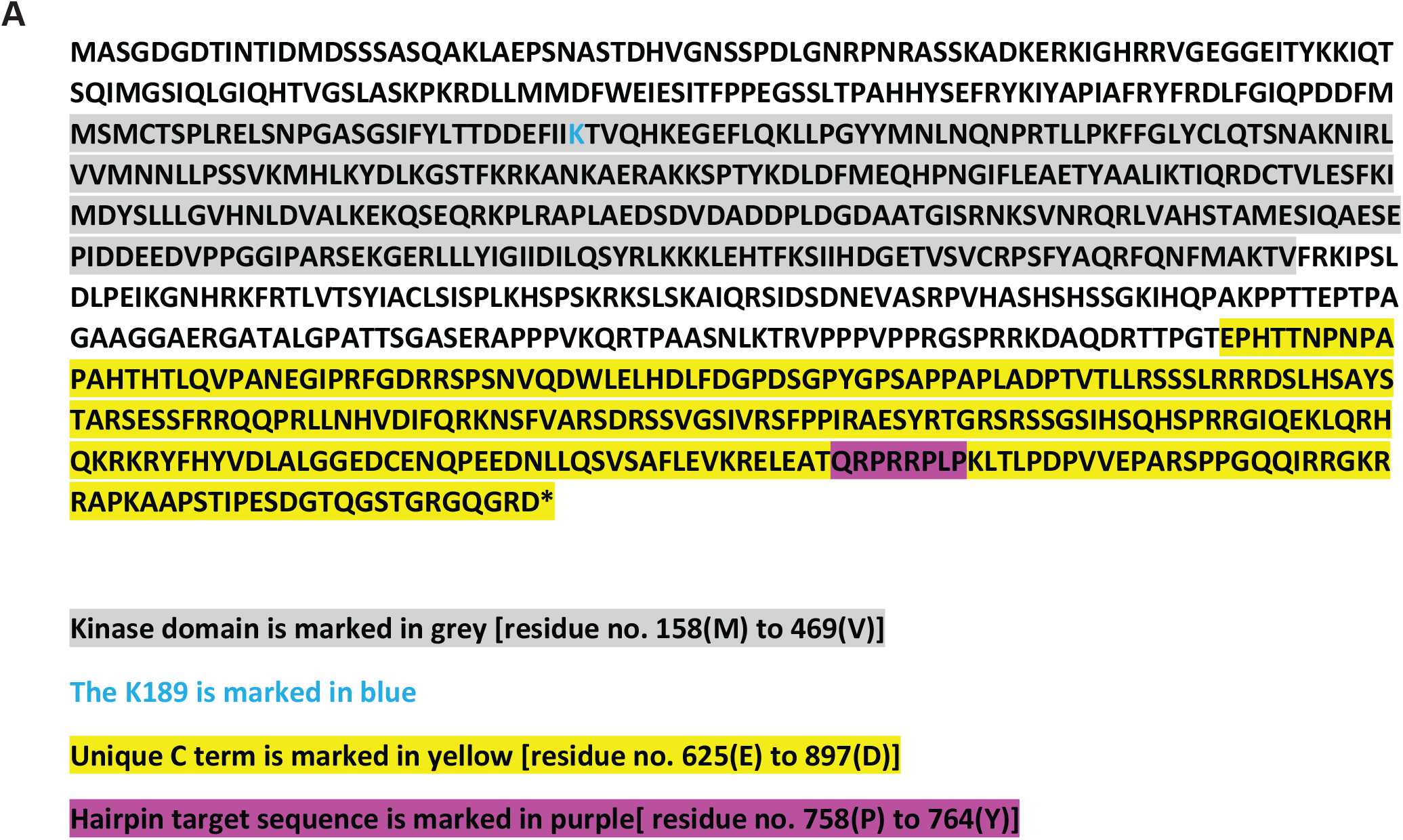
**(A)** Full protein sequence of dPIP5K^L^. The kinase domain is highlighted in grey and the unique C terminal region of dPIP5K^L^ is marked in yellow. The K189, used to make point mutant to generate kinase dead flies is highlighted in blue. Purple marks the residues targeted by hairpin construct against the C terminal of dPIP5K^L^.

**Supplementary Fig S2:**
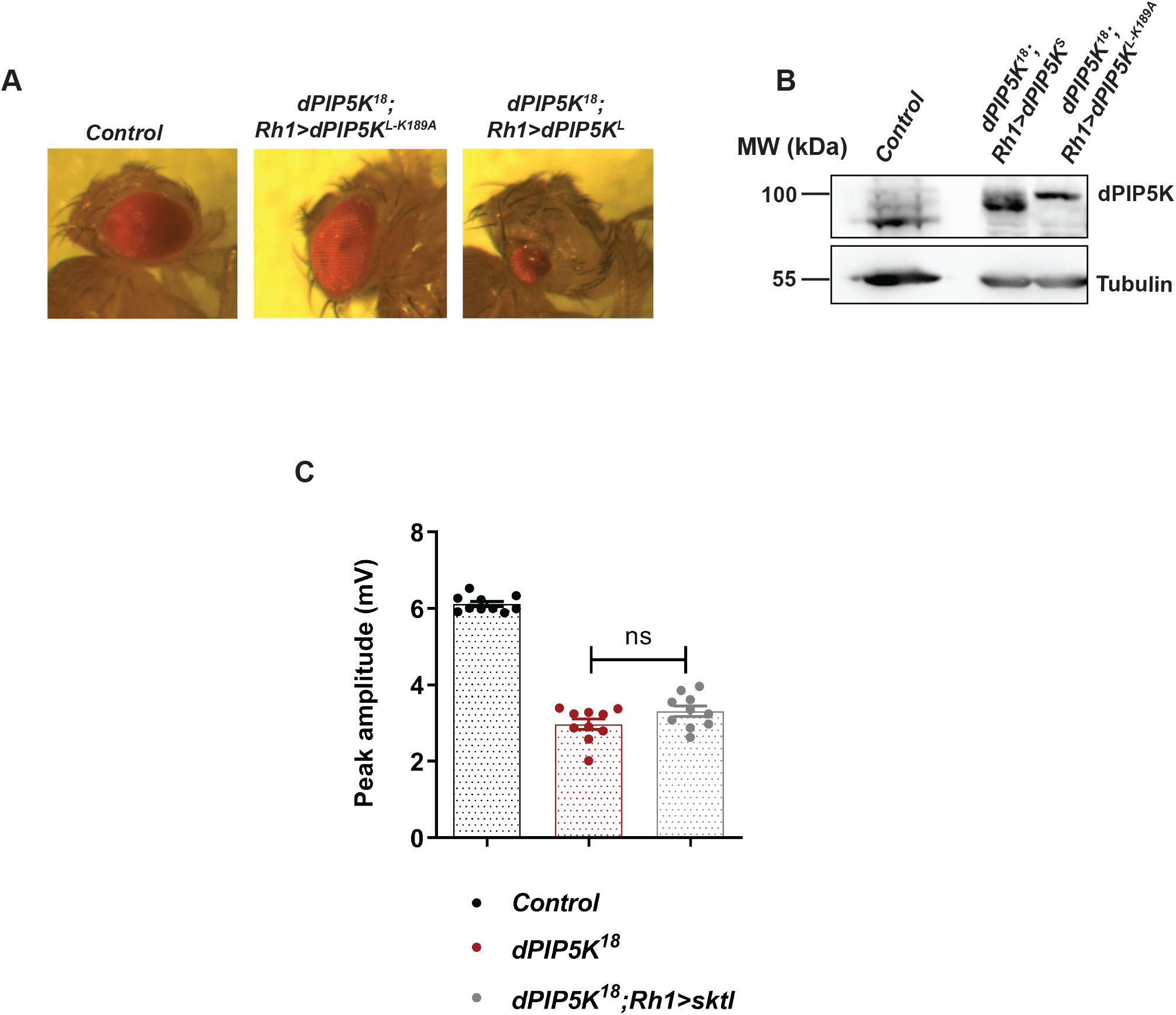
**(A)** Images showing eye morphology of freshly eclosed flies of the mentioned genotypes. The full-sized red eye of control flies is shown; eye with morphological defect seen on overexpressing dPIP5K^L^ but not upon over expression of dPIP5K^L-K189A^. **(B)** Western blot from head extracts of 1- day old fly of the mentioned genotypes showing expression of either dPIP5K^S^ or dPIP5K^L-K189A^ (100kDa) in dPIP5K^18^ background. Tubulin at 55kDa is used as loading control **(C)** Quantification of the peak amplitude for ERG response in each of the mentioned genotypes. Y axis represents peak amplitude and X axis shows the genotypes. Each data point represents an individual fly tested. (n=10 flies). Error bars represent S.E.M. Data information: Scatter dot plot with mean±S.E.M is shown. Statistical tests: (C) One way Anova with post hoc Tukey’s multiple pairwise comparison. ns-Not significant

**Supplementary Fig S3:**
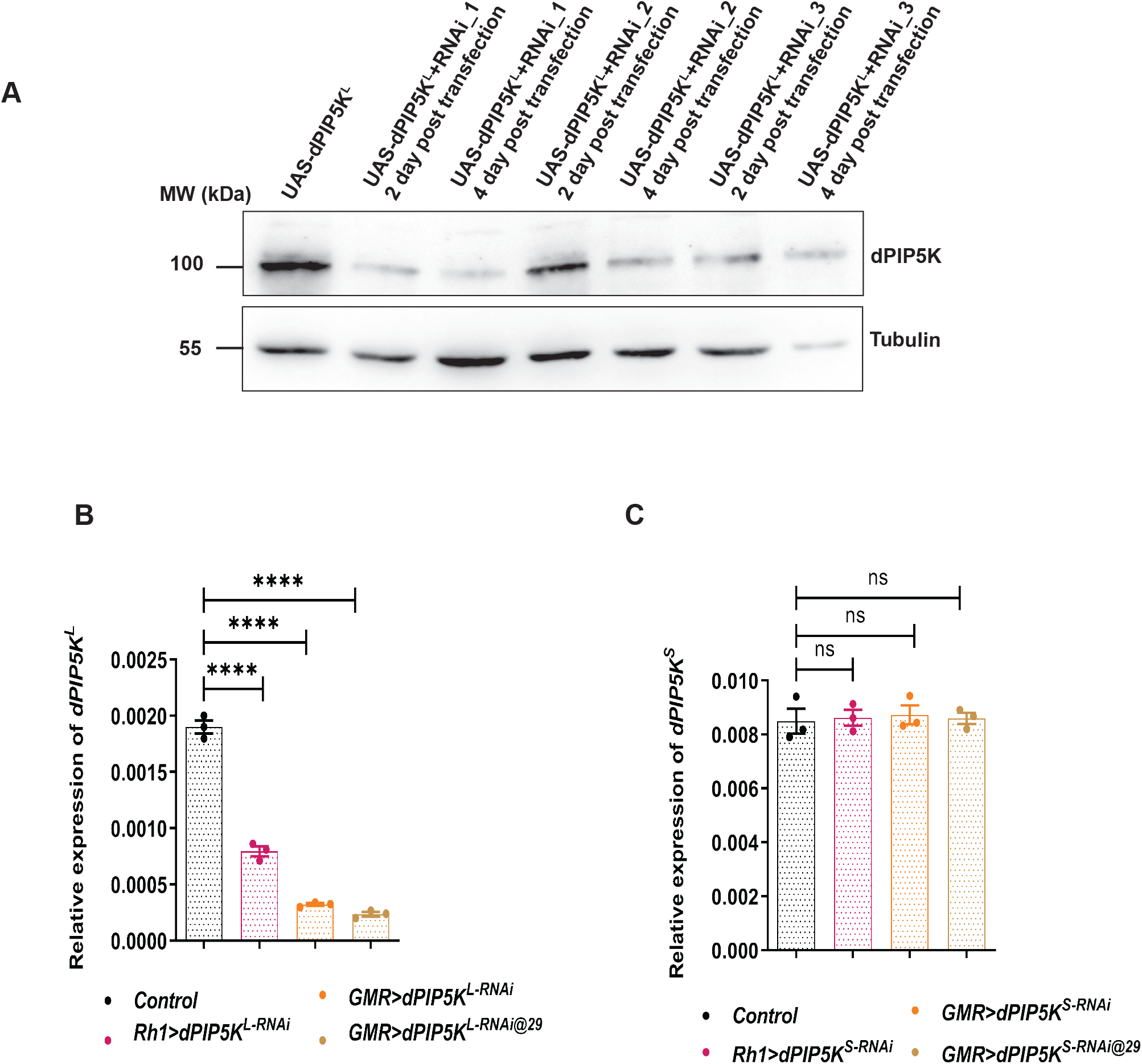
**(A)** Western blot from S2R+ cell extract upon co-transfection with *dPIP5K^L^* and hairpin constructs against the C terminal region of dPIP5K^L^ is shown. dPIP5K^L^ is detected at 100 kDa and Tubulin at 55kDa is used as loading control. Construct 1(RNAi_1) gives the maximum knockdown. **(B)** Quantitative real time PCR from retinae of 1- day old flies showing the degree of *dPIP5K^L^* transcript knockdown for the mentioned genotypes. Y axis represents relative transcript level for *dPIP5K^L^*, normalized to internal control (*RP49*: ribosomal protein 49). X axis represents the genotypes (n=3 biological replicates). Error bars represent S.E.M. **(C)** Quantitative real time PCR from retinae of 1- day old flies showing no change of transcript level of *dPIP5K^S^* upon knockdown of dPIP5K^L^ in the mentioned genotypes. Y axis represents relative transcript level for *dPIP5K^S^*, normalized to internal control (*RP49*: ribosomal protein 49). X axis represents the genotypes (n=3 biological replicates). Error bars represent S.E.M. Data information: Scatter dot plots with mean±S.E.M are shown. Statistical tests: (C) One way Anova with post hoc Tukey’s multiple pairwise comparison. ns-Not significant; ****p<0.0001

